# SIV and *Mycobacterium tuberculosis* synergy within the granuloma accelerates the reactivation pattern of latent tuberculosis

**DOI:** 10.1101/2020.02.21.959353

**Authors:** Collin R Diedrich, Tara Rutledge, Pauline Maiello, Tonilynn M Baranowski, Alexander G White, H. Jacob Borish, Paul Karell, Forrest Hopkins, Jessica Brown, Sarah M Fortune, JoAnne L Flynn, Zandrea Ambrose, Philana Ling Lin

## Abstract

Human immunodeficiency virus infection is the most common risk factor for severe forms of tuberculosis (TB), regardless of CD4 T cell count. Using a well-characterized cynomolgus macaque model of human TB, we compared radiographic, immunologic and microbiologic characteristics of early (subclinical) reactivation of latent *M. tuberculosis (*Mtb) infection among animals subsequently infected with simian immunodeficiency virus (SIV) or who underwent anti-CD4 depletion by a depletion antibody. CD4 depleted animals had significantly fewer CD4 T cells within granulomas compared to Mtb/SIV co-infected and Mtb-only control animals. After 2 months of treatment, subclinical reactivation occurred at similar rates among CD4 depleted (5 of 7 animals) and SIV infected animals (4 of 8 animals). However, SIV-induced reactivation was associated with more dissemination of lung granulomas that were permissive to Mtb growth resulting in greater bacterial burden within granulomas compared to CD4 depleted reactivators. Granulomas from Mtb/SIV animals displayed a more robust T cell activation profile (IFN-α, IFN-γ, TNF, IL-17, IL-2, IL-10, IL-4 and granzyme B) compared to Mtb/αCD4 animals and controls though these effectors did not protect against reactivation or dissemination, but instead may be related to increased viral and/or Mtb antigens. SIV replication within the granuloma was associated with reactivation, greater overall Mtb growth and Mtb killing resulting in greater overall Mtb burden. These data support that SIV disrupts protective immune responses against latent Mtb infection beyond the loss of CD4 T cells, and that synergy between SIV and Mtb occurs within granulomas.

**Author Summary:** Most humans are able to control infection with *Mycobacterium tuberculosis* (Mtb), the bacteria that causes tuberculosis (TB). Controlled, asymptomatic infection (latent infection) can develop into symptomatic, severe TB (reactivation TB) when the immune system is impaired, and HIV is the most common risk factor. Chronic HIV infection is associated with low CD4 T cells but there are likely other factors involved. Using macaques with latent Mtb infection, we could induce reactivation from either CD4 T cell depletion or SIV infection. We found that SIV induced reactivation was more dramatic with more bacterial dissemination and bacterial growth compared to those with CD4 depletion. While SIV-infected animals had more activated immune responses in the lung granulomas (a collection of immune cells that functions to combat Mtb), they could not prevent bacterial spread of Mtb resulting in more TB pathology, higher bacterial burden and disease throughout the body. These data suggest that the HIV-induced reactivation TB is not solely from the loss of CD4 T cells. Furthermore, our data suggest that SIV and Mtb have a synergistic relationship within granulomas that impairs the ability to kill Mtb and that this relationship exacerbates TB disease.

## Introduction

Tuberculosis (TB) continues to be a major health concern, with an estimated 10 million new cases of TB in 2018. Of the 1.5 million TB deaths that year, an estimated 251,000 were in human immunodeficiency virus (HIV)-infected individuals [1]. The majority (∼90%) of immune competent individuals infected with *Mycobacterium tuberculosis* (Mtb) develop an asymptomatic state of controlled infection called latent infection (LTBI), while others develop symptomatic or active TB [1]. HIV infection increases host susceptibility to TB [2] and pathology [3], including primary TB (symptomatic TB that develops soon after Mtb infection) or reactivation of LTBI. HIV-infected individuals are approximately 9 times more likely to develop reactivation from LTBI than HIV-uninfected individuals [4]. TB incidence in HIV+ persons increases as peripheral CD4 T cell numbers decline, suggesting that CD4 T cells are important in control of Mtb infection [2], which is supported by animal model studies [5]. However, HIV-infected individuals with normal peripheral CD4 T cell counts are still more susceptible to active TB than their HIV-uninfected counterparts [2, 3]. This leads to the hypothesis that the HIV-associated increase in TB susceptibility is not solely due to the loss of CD4 T cells.

The histopathologic hallmark of TB is the granuloma. Granulomas are organized immunological structures composed of T cells, macrophages, B cells, NK cells, dendritic cells and other immune cells that surround Mtb to form both a physical and immunologic barrier to prevent Mtb dissemination. As a respiratory infection, these granulomas are most prominent in the lung, but can also be present in the mediastinal lymph nodes and other organs (reviewed in [6]). While granulomas can kill Mtb under optimal immune conditions, they can also be a site for bacterial persistence and/or growth particularly during latent infection. Cynomolgus macaques infected with low dose Mtb develop the full spectrum of human infection outcomes, from latent to active TB, with histopathologic features of granulomas nearly identical to human [7, 8]. From this model, we have learned that the immune factors within each granuloma are variable and complex, reflecting a delicate balance between pro- and anti-inflammatory cytokines necessary for optimal function [9].

Human studies that examine *M. tuberculosis* granulomas within HIV-coinfected individuals are informative but highly variable [10], necessitating non-human primate (NHP) models to understand how *M. tuberculosis* granulomas change during co-infection. NHP are an invaluable animal model to study SIV and *Mtb* co-infection [11–17]. Use of these models facilitates a more in-depth understanding of how pre-existing infection can influence the outcome of co-infection and the immunologic mechanisms of worsening disease. In human co-infection, it is not generally known which infection occurred first (HIV or Mtb) or the duration of each infection before subjects come to clinical attention. SIV infection prior to *Mtb* infection was associated with increased acute TB pathology with extrapulmonary dissemination and increased bacterial burden [13, 17]. In contrast, NHPs with established latent Mtb infection and subsequent SIV infection had variable rates of reactivation TB depending on the time point after SIV infection [11, 14, 15]. These studies of SIV-induced reactivation of LTBI have suggested that loss of CD4 T cells can contribute to reactivation of LTBI but likely CD4 T cell independent mechanisms are important as well [11, 12, 15, 16]. In our previous studies, SIV infection of cynomolgus macaques with LTBI induced reactivation in all animals [11, 12], with some reactivating early after SIV infection while others did not reactivate until up to 10 months after SIV infection. Early reactivation was associated with greater peripheral CD4 T cell depletion within the first 8 weeks suggesting that CD4 T cells played an important role in one aspect of reactivation but not in all cases [11, 12]. Similarly, latently Mtb-infected NHP given humanized CD4 depletion antibody had a 50% reactivation rate [5], where reactivation was associated with more severe depletion of CD4 T cells in the mediastinal lymph nodes and not with extent of peripheral CD4 depletion [5]. These studies and those of others [11, 14, 15] have suggested that granuloma specific responses (including resident CD4 T cell function) are more critical than peripheral CD4 T cell counts.

The advent of more sophisticated tools to assess pathogenesis, bacterial dissemination and disease progression have improved our understanding of how LTBI is established and the events that result in reactivation. Using positron emission tomography and computed tomography (PET CT), we have shown that a variety of patterns exist in clinically defined LTBI and this spectrum of latency influences the risk of tumor necrosis factor (TNF)-neutralization-induced reactivation [18]. In that study, the risk of reactivation was associated with specific PET CT characteristics including overall lung inflammation and individual characteristics of lung granulomas [18]. This latter finding is consistent with previous published data that granulomas are independent from each other with their own bacterial burden and immune response [9, 19]. HIV-infected humans with LTBI who had PET CT-identified subclinical TB disease were more likely to develop clinical disease than those without subclinical pathology [20], demonstrating the similarities between humans and this NHP model.

In prior studies, reactivation of LTBI was defined by NHP displaying overt signs of disease (e.g., coughing, weight loss, respiratory distress, abnormal X-ray) [5, 11, 12]. Here, we extend our previous studies to compare the radiologic, immunologic, and microbiologic characteristics during the earliest phases of reactivation TB before overwhelming disease pathology and bacterial burden occurs, which could potentially bias interpretation of immune data. We previously established the rates of reactivation after LTBI in both CD4 depleted and SIV_mac251_ infected animals and therefore could predict when early, subclinical reactivation would begin. Moreover, we also established a method to detect subclinical reactivation where the formation of new granulomas detected by PET CT during established LTBI indicated a disruption within the host immune response before NHP showed signs of overt clinical reactivation [18]. Thus, NHP with established LTBI underwent SIV infection (Mtb/SIV), αCD4 depletion antibody treatment (Mtb/αCD4), or no immune suppression (Mtb-only, controls) and were serially assessed by PET CT with granuloma specific bacterial burden, SIV replication, pathology, immunology, and disease progression compared at necropsy. Despite having more CD4 T cells than Mtb/αCD4 macaques, Mtb/SIV-coinfected animals had greater dissemination of lung granulomas observed by PET CT and confirmed at necropsy that were more permissive to Mtb growth. SIV replication within individual granulomas was associated with reactivation occurrence, reduced Mtb killing and increased Mtb growth. The frequency and functional profiles of T cells within granulomas differed significantly between Mtb/SIV and Mtb/αCD4 groups during subclinical reactivation. These data support that SIV infection has multiple mechanisms of disrupting the protective immune response against Mtb that are independent of CD4 depletion, and that SIV exerts local effects on the immune response and Mtb within individual granulomas highlighting the synergy between SIV and Mtb within individual granulomas.

## Methods

### Animals

Adult (> 4 years of age) cynomolgus macaques (*Macaca fascicularis*) were screened for other co-morbidities (e.g., parasites, SIV, Mtb) before challenge (Valley Biosystems, Sacramento, CA). Animals were infected with low dose (∼ 15 CFU per monkey) of *M. tuberculosis* (barcoded Erdman strain [21]) via bronchoscopic instillation to the lower lung lobe and housed in Biosafety Level 3 (BSL-3) NHP facility. Cynomolgus macaques inoculated with Erdman *M. tuberculosis* was used in this study because latent and active *M. tuberculosis* infection has been extensively characterized [5, 7–9, 11, 12, 18, 19, 21–24]. Mtb infection was confirmed by the detection of TB-specific lesions on serial PET CT scans and identified at necropsy. As in prior studies in this LTBI model, asymptomatic animals with no culturable Mtb in bronchoalveolar lavage (BAL) or gastric aspirate samples, and normal erythrocyte sedimentation rate (ESR, marker of systemic inflammation) were declared with LTBI at 6 months post-Mtb infection, similar to human clinical definitions [7]. Animals that developed active TB were excluded and moved to a different study. After latent Mtb infection was established, animals were randomized to receive either intravenous challenge with a viral swarm SIV_mac251_ (1.67×10^5^ viral RNA copies) [11, 12] (n=8), CD4 depletion (rhesus recombinant depleting CD4 antibody, 50mg/kg/dose IV every 2 weeks until necropsy [5]) (n=7), saline (Mtb-only control, n=6) for 8 weeks. Stratification into treatment groups was based on the total lung FDG activity by PET CT to ensure that there was no potential bias toward reactivation within any experimental group as increased total lung FDG activity was associated with increased risk of reactivation during TNF neutralization [18]. Four macaques were infected with SIV_mac251_ only for 8 weeks as a SIV-only control group.

Blood was obtained via venipuncture for isolation of peripheral blood mononuclear cells (PBMC) every 1-4 weeks as previously described [18]. Bronchoalveolar lavage (BAL) was performed and peripheral (axillary or inguinal) lymph nodes (pLN) were biopsied to measure tissue specific CD4 depletion in both SIV and CD4 depletion groups at serial time points (0, 3 and 7 weeks after SIV infection or CD4 depletion), as previously described [11].

### PET-CT imaging and analysis

*In vivo* disease progression was assessed using PET co-registered with CT with a microPET Focus 220 preclinical PET scanner (Siemens Medical Solutions) and clinical 8 slice helical CT scanner (NeuroLogica Corp) as previously described [24, 25]. The PET probe used was 2-deoxy-2-^18^F-D-deoxyglucose (FDG) as we have previously shown that this can be used to identify TB lesions [24]. PET CT scans were performed every four weeks after Mtb infection until 6 months post-infection when latent Mtb infection was declared. Prior to SIV infection or CD4 depletion, a scan was performed to determine baseline disease and then every 2 weeks until the time of necropsy (8 weeks later or earlier if signs of clinical deterioration developed). Degree of Mtb involvement within the lungs and mediastinal lymph nodes was measured using several different parameters as previously described [25] which included: identification and count of individual granulomas, total lung FDG activity, single granuloma FDG avidity and size over time, number of mediastinal lymph nodes with increased FDG avidity with or without the presence of necrosis, presence of extrapulmonary involvement (e.g., liver lesions). Each PET CT scan was analyzed by a team of blinded analysts (PM, AGW, HJB).

### Subclinical reactivation of latent *M. tuberculosis* infection by PET CT

We previously showed that SIV_mac251_ infection caused clinical reactivation in 100% of our latently infected NHP (defined by signs of disease such as coughing or weight loss, new growth of *M. tuberculosis* by BAL and gastric aspirates, abnormal X-ray, and increased ESR) between 12-48 weeks post SIV infection [11]. The goal of this study was to identify the pathologic, microbiologic, and immunologic changes during subclinical reactivation. To do this, PET CT was used to define reactivation as the formation of a new granuloma (whose presence was confirmed at necropsy) after latent *M. tuberculosis* infection was established [18]. This more stringent definition of reactivation allows us to identify these changes before overwhelming clinical signs of disease developed and immune responses could be completely confounded by profoundly high bacterial burden and pathology.

### Necropsy

At necropsy, Mtb-involved tissue sites (i.e. individual granulomas and other pathologies) were matched by PET CT and harvested for analysis as previously described [17-19, 21, 26]. Gross pathology of TB was quantified by assessing the number, size, and pattern of granulomas distributed within each lung lobe, thoracic lymph nodes and in extrapulmonary sites as previously published [22]. PET CT matched individual granulomas and other tissues harvested at necropsy were homogenized into single cell suspension and plated for Mtb and analyzed by multiparameter flow cytometry [9].

### *Ex vivo* immunologic assays

Peripheral blood CD4 and CD8 T cell measurements were obtained every 2 weeks after SIV and CD4 depletion. Mtb specific immune responses from tissues were measured by stimulating with Mtb ESAT6-CFP10 peptide pools (10µg/ml of each peptide) or incubated with only media (RPMI+10%hAB). Cells from tissues at necropsy were isolated and stimulated for 4 hours with peptide pools in the presence of Brefeldin A.

Flow cytometry was performed on all isolated cells after stimulation by staining with a combination of antibodies, as described [9]. PBMC and BAL were stained for CD4 and CD8 T cells (CD3, CD4, CD8). Granuloma cells were stained for CD3, CD4, CD8, granzyme B, IL-4, IFN-γ, IL-2, IL-10, IL-17, and TNF. Data acquisition was performed using an LSR II (BD) and analyzed using FlowJo Software v.9.7 (Treestar Inc, Ashland, OR). Only lung granulomas with more than 40 lymphocytes (median: 397, IQR_25-75_: 121-1545) were included in cell frequency analyses to ensure all granulomas contained an accurate estimate. When measuring cytokine production and granzyme B presence, only samples with a minimum of 100 CD3 T cells by flow cytometry (median: 631, IQR_25-75_: 205-14919) were included to ensure precise and accurate measurements, as published [9, 27]. Gating strategies used for analysis are presented using ESAT6-CFP10 stimulated cells from the granuloma (Supplemental Figure 1).

### Viral RNA Quantification

Plasma CD4 and SIV viral RNA copies from each designated timepoint were performed in bulk from filtered (0.45 um to remove any potential Mtb) plasma samples. RNA extractions from pelleted plasma were performed by QiAmp viral miniRNA protocol (Qiagen, Venlo Netherlands) per the package insert instructions. Tissue specific RNA extraction was performed by first homogenizing tissues into single cell suspension and freezing samples in a 1:4 ratio of homogenate to Trizol LS (Thermo Fisher Scientific, Waltham Massachusetts) and stored (−80°C). Tissue RNA extraction was performed using the Direct-zol RNA miniprep plus kit with DNAase treatment per package insert (Zymo, Irvine, California). SIV and CD4 primers for RT PCR and amplification conditions were as previously published [28]. RT-PCR was performed using the QuantStudio 6 Real-Time PCR system (Thermo Fisher Scientific, Waltham Massachusetts).

### Mtb bacterial burden estimates, genome isolation, quantification, Mtb killing and barcode mapping

Mtb burden was estimated by colony forming units (CFU) of single cell homogenate from each individual site, as previously described [18, 22]. Lymph node burden (“LN CFU”) was defined as the sum of CFU from all mediastinal lymph nodes. Extrapulmonary score (EP score) is a quantitative estimate of extrapulmonary involvement (e.g., liver, peripancreatic lymph node, paracostal abscess, kidney) for which bacterial growth, gross or microscopic evidence of Mtb involvement are taken into account [22]. Total bacterial burden includes the sum of CFU from the lymph nodes (mediastinal and extrapulmonary) and lung lesions (e.g., grossly normal lung, granulomas, involved lung, or diaphragm granulomas).

Mtb DNA extractions and qPCR for estimating chromosomal equivalents were performed as previously described [23]. Chromosomal equivalents (CEQ) were assessed relative to a serially diluted standard curve of *M. tuberculosis* genomic DNA using quantitative real-time PCR; efficiency for each run was kept between 90% and 110%. Each sample was analyzed in triplicate on a ViiA7 real-time PCR system (ThermoFisher Scientific, Waltham Massachusetts) with a 384-well block Quantification of CEQ using primers targeting SigF and iTaq Universal SYBR Green Supermix (Bio-Rad, Hercules, CA).

Genetically barcoded Mtb was designed by inserting random identifier tags into the Mtb chromosome as already published [21]. We previously showed that each granuloma is established by a single individual Mtb bacillus [19]; and therefore are able to use the digitally barcoded Mtb to facilitate mapping of bacterial dissemination. Mtb genomes were extracted and sequenced from Mtb colonies grown from individual tissues (e.g., granulomas, lymph nodes) harvested at necropsy. Individual barcode identities were determined by Mtb genome sequences via a customized pipeline [21] and each barcode was matched with the 3-dimensional x,y,z coordinates of the lungs on PET CT. Barcodes from granulomas observed prior to immune suppression and new granulomas that appeared after immune suppression were compared.

### Immunohistochemistry

Immunohistochemistry was performed as previously described [23]. A portion of each lung granuloma was formalin-fixed and paraffin embedded. After deparaffinization of processed slides, antigen retrieval was performed using boiling EDTA Tris (pH 9) buffer under pressure for 6 minutes. Tissues were blocked using 1% bovine serum albumin (BSA) and stained for CD3 (1:200, monoclonal rat Ab11089; Abcam Cambridge, MA) and CD38 (1:1000 polyclonal rabbit A9696, Lifespan Biosciences, Seattle WA) overnight at 4°C. Appropriate florescent secondary antibodies were used to visualize primary antibodies or as secondary isotype controls. Dapi was utilized to identify nuclei. Enumeration of CD38 and CD3 co-localization was manually counted. Images were acquired using a Nikon 90I epi-fluorescent microscope (Nikon, Melville, NY) at 20x objective with Nikon Elements AR 4.51.00 64-bit.

### Statistics

Shapiro-Wilk test was used to test for normality. Treatment groups were compared using a Wilcoxon-exact test (also known as the Mann-Whitney test) (for 2 group comparison) or a Kruskal-Wallis with Dunn’s multiple comparison adjusted p-values reported (for 3 or more group comparisons) or Steel multiple comparison adjustment (for 2 comparisons). The Pearson correlation coefficient and corresponding p-values were reported for relationships among normally-distributed variables and the Spearman correlation coefficient was reported for nonparametric data. All statistical tests on serial data were performed in JMP Pro 14.0.0 (64-bit, SAS institute, Cary, NC). For group comparisons on non-serial data, Graphpad Prism Mac OSX (Version 8.2.1, GraphPad San Diego, CA) was used. For counts (including cell counts, CFU, and FDG activity), the data was first transformed (adding 1 to entire dataset), so that zeroes could be visualized and analyzed on a log_10_ scale. All statistical tests are two-sided and significance was established at p ≤ 0.05. Permutations tests for comparison of T cells proportions, presented in pie charts, was performed using SPICE 6.0 (NIH, Bethesda MD) [29].

A principal components analysis was performed (using JMP Pro) on the cytokine expression from absolute numbers of both CD4 and CD8 cell counts (log_10_ transformed) from individual granulomas. Using the Kaiser criterion (dropping any components with eigenvalues less than 1), the first principal component was saved as a new variable for both CD4 and CD8 cell types.

To ensure against bias from any single animal in our lung granuloma data, the median frequency and absolute counts of CD4 and CD8 of each animal were calculated and effect sizes between groups examined. We found similar effect size among the CD4 T cells frequency and absolute CD8 T cells. The CD8 T cell effect size between SIV and control was inflated due to one animal (31316) but the overall trends were similar. A similar comparison was performed with the cytokine data used in the principal component analysis with similar effect sizes.

### Ethics statement

All animal protocols and procedures were approved by the University of Pittsburgh’s Institutional Animal Care and Use Committee (IACUC) that adheres to the national guidelines established in the Animal Welfare Act and Guide for the Care and Use of Laboratory Animals as mandated by the U. S. Public Health Service Policy (PHS). The IACUC approval number for our study is 17050656 and our PHS policy number is D16-00118.

University of Pittsburgh housed all NHP in temperature, humidity, and lighting controlled rooms. Single housed cages at least 2 square meters apart were utilized, allowing for visual and tactile contact with neighboring NHP. NHP were fed twice daily with specific formulated biscuits and at least 4 days/week with fruits and vegetables and had *a libitem* access to water. An enhanced enrichment plan was designed and administrated by NHP enrichment specialists. Species-specific behaviors were always encouraged. NHP maintained constant access to toys and other manipulata. All manipulata and toys were regularly rotated. Puzzle feeders and cardboard tubing were used to simulate foraging for food and adjustable mirrors were utilized to simulate interactions with other animals. Regular human and NHP interactions were encouraged. These interactions consisted of administering small food objects that follow all safety protocols. Caretakers interact with NHP by talking or use of non-aggressive facial expressions while performing housing area tasks. All feedings, cage cleanings, and other routine procedures were completed on a strict schedule to allow NHP to acclimate to a normal daily schedule. All NHP were provided a diverse variety of visual and auditory stimulation, which included either radios or video equipment that are designed for children for at least three hours a day. These radios and video were rotated between animal rooms to avoid too much repetition for the same housed animals.

Appetite, attitude, activity level, hydration status, etc. were documented two times daily to ensure the health of each NHP. After Mtb infection, NHP were monitored for signs of TB disease (e.g., anorexia, weight loss, tachypnea, dyspnea, coughing). Physical exams were performed on a regular basis, as well. NHP were sedated prior to any veterinary procedure using ketamine or other approved drugs. Veterinary technicians regularly document disease progression through regular PET CT imaging and closely monitor all NHP for signs of pain or distress. If any signs of pain or distress are identified appropriate supportive care (e.g. dietary supplementation, rehydration) and clinical treatments (analgesics) are given. If any NHP has advanced disease or intractable pain they are sedated with ketamine and then humanely euthanized using a lethal dose of sodium pentobarbital.

## Results

### αCD4 antibody results in more dramatic and sustained CD4 reduction than SIV infection

Latently infected NHP were stratified to control, αCD4 depleting antibody or SIV infection groups. SIV infection was confirmed by quantitative RT-PCR of viral RNA in plasma (Figure 1A). SIV induced a transient reduction in peripheral CD4 T cells that was most dramatic 3 weeks after infection, which coincided with peak viral RNA copies (Figure 1B), as previously observed for this viral strain [11]. With the exception of this time point (3 weeks post SIV infection), CD4 and CD8 T cell frequencies and absolute counts in the blood were similar to Mtb-only control groups. The CD4 T cells in the Mtb/SIV NHP BAL were transiently lower than LTBI controls but not in the peripheral lymph nodes (pLN). CD4 depletion antibody caused a significant reduction in frequency and absolute numbers of CD4 T cells from baseline across multiple time points in blood (Figure 1B), airways (Figure 1C) and within peripheral lymph nodes (Figure 1D) as reported previously [5]. Overall, latently infected NHP undergoing CD4 depletion (Mtb/αCD4 NHP) had more severe and sustained reductions in CD4 T cells in the blood and pLN compared to animals undergoing SIV infection (Mtb/SIV NHP) and LTBI control groups.

**Figure 1.**
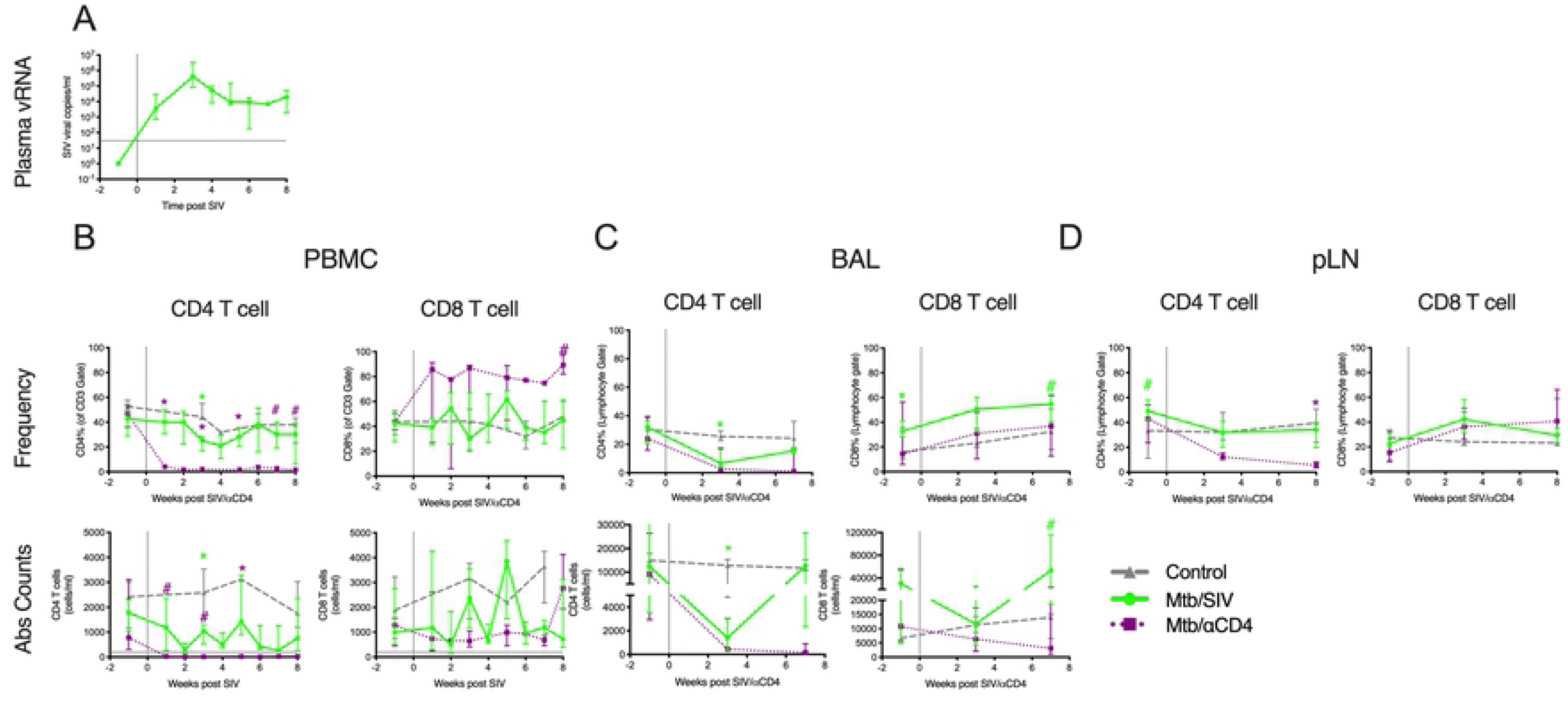
Changes in plasma viral RNA copies and T cells in peripheral blood mononuclear cells (PBMC), bronchoalveolar lavage (BAL), and peripheral lymph node (pLN) over time. A) Plasma viral RNA copies among Mtb/SIV co-infected animals are shown. B) Peripheral CD4 T cells are more severely reduced compared to Mtb/SIV co-infected animals with latent infection. Plasma viral RNA is reported as mean and standard deviation. C) Total absolute CD4 and CD8 T cell counts and frequencies were measured in BAL cells. D) CD4 and CD8 T cell frequencies were measured in pLN. Changes in absolute CD4 and CD8 T cell counts (Abs Counts) and frequencies after SIV_mac251_ infection (green line, Mtb/SIV, n = 8), αCD4 depletion antibody (purple line, Mtb/αCD4, n = 7) and controls (grey line, Mtb-only control, n = 6) are shown. Statistics reported are Steel tests comparing Mtb/SIV and Mtb/αCD4 at each time point (adjusted for comparing Mtb-only controls to Mtb/SIV [green stats marker] and Mtb/αCD4 to Mtb/SIV [purple stats marker]). For B-D, medians are shown with error bars representing interquartile range. * p < 0.05, # p < 0.10.

### Reactivation characteristics differ between CD4 depletion and Mtb/SIV infected animals

We previously published that both CD4 depletion [5] and SIV_mac251_ infection [11] during LTBI resulted in reactivation rates of 50% (within 12 weeks) and 100% (50% by 8 weeks and 100% by 48 weeks after SIV), respectively. To better characterize the pathogenesis of reactivation due to these interventions without profoundly perturbing the immune response with overwhelming bacterial burden and severe pathology, subclinical reactivation was used as an endpoint. Subclinical reactivation of LTBI was defined by the appearance of a new lung granuloma by PET CT after immune suppression (representing Mtb dissemination and impaired immune control), as previously published [18]. The presence of these new granulomas observed by PET CT was confirmed by gross pathology at necropsy.

Despite a greater reduction of CD4 T cells by αCD4 depletion antibody than SIV_mac251_ (Figure 1), the strict definition for subclinical reactivation resulted in similar rates of reactivation in Mtb/SIV (4 of 8 or 50%) and Mtb/αCD4 (5 of 7 or 71%) animals during the 8 weeks of treatment after LTBI (Supplemental Table 2). Because this study was powered based on each experimental group against LTBI control and not by reactivation status within each group, statistical power to examine rates of reactivation between groups was limited. Among animals with reactivation, only 1 of the 5 animals in the Mtb/αCD4 group had clinical signs (i.e., increased respiratory effort) compared to all 4 of the Mtb/SIV animals with reactivation (i.e., lethargy, increased respiratory effort). All 5 of the reactivation animals in the Mtb/αCD4 group and all 4 in the Mtb/SIV group had elevated ESRs (a systemic marker of inflammation), including 4 animals that also had growth of Mtb detected by gastric aspirate (GA) or BAL (2 in each group, Supplemental Table 2). At necropsy, the degree of TB-specific gross pathology (determined by necropsy score [22]) was similar between reactivators and non-reactivators receiving CD4 depletion (Figure 2A), while a trend (p = 0.096) toward higher necropsy score was observed among reactivators of the Mtb/SIV group. Some of the scores may have been underestimated if animals were euthanized early (i.e., one Mtb/αCD4 NHP suffered an unrelated aneurysm requiring early necropsy, one of the Mtb/SIV NHP developed overt PET CT signs of reactivation, and one Mtb/SIV NHP developed clinical signs of deterioration). Both reactivators and non-reactivators in the Mtb/αCD4 group had similar Mtb burden in the lungs and lymph nodes (Figure 2B) with similar extrapulmonary involvement (Figure 2A). In contrast, Mtb/SIV reactivated animals had greater total bacterial burden and lung burden compared to non-reactivators (Fig 2B). A greater proportion of granulomas with Mtb growth was observed among Mtb/SIV reactivators compared to non-reactivators (Figure 2C). All Mtb/SIV reactivated animals had extrapulmonary involvement (Fig. 2A). A positive correlation was observed between Mtb growth within thoracic lymph nodes and extrapulmonary involvement among Mtb/SIV NHP but not in the control or Mtb/αCD4 NHP (Supplemental Figure 2). Limited bacterial killing in the lymph nodes has been previously observed during Mtb infection, with lymph node involvement positively correlated with extrapulmonary disease [23]. These data suggest that lymph nodes may be a source of bacterial dissemination [23]. Thus, despite higher CD4 T cell levels in the SIV infected groups compared to Mtb/αCD4 NHP, SIV infection induced more dramatic changes in disease and bacterial burden than CD4 depletion. Furthermore, the changes that occur during early, subclinical CD4 depletion induced reactivation are more subtle than SIV-induced reactivation and are not easily detected by our current gross pathology metrics (i.e., necropsy score) of TB disease at necropsy.

**Figure 2.**
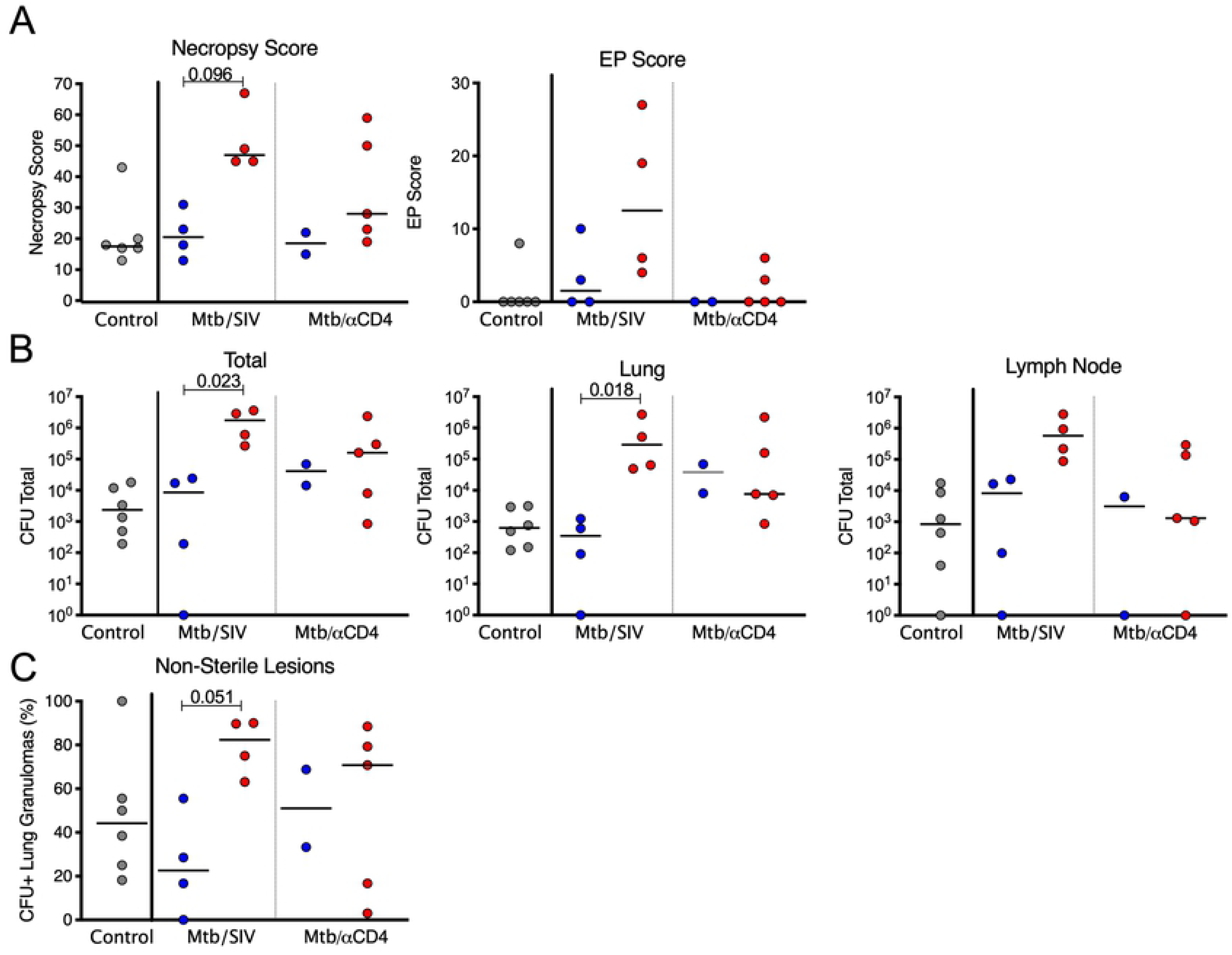
Subclinical reactivation of Mtb/SIV NHP results in greater total thoracic burden but not in Mtb/αCD4 NHP. Non-reactivators (blue) and reactivators (red) from Mtb only (control, grey), Mtb/SIV co-infected (n = 8), and CD4 depletion (Mtb/αCD4, n = 7) NHP are shown. A) Necropsy and extrapulmonary (EP) scores are based on gross pathology at time of necropsy. B) Total thoracic burden (quantitative sum of Mtb from excised tissues within the thoracic cavity) and lung and thoracic lymph nodes are shown. C) A greater percentage of granulomas with Mtb growth is observed in reactivated Mtb/SIV NHP. P-values reported from Kruskal-Wallis test with Dunn’s multiple comparisons adjustments, adjusted for the following (4) comparisons: reactivators vs non-reactivators within treatments and non-reactivators and reactivators between treatments. P-values < 0.10 are shown. Each dot represents an animal. Mtb-only controls are shown for reference, but not included in the statistical analysis.

We sought to further characterize the differences in reactivation patterns between the two groups. Reactivators in the Mtb/SIV group had more new granulomas (median = 19.5 new granulomas per NHP) observed by PET CT compared to the Mtb/αCD4 group (median= 2 new granulomas per NHP), though it was not statistically significant given the heterogeneity that is inherent within this NHP model (Figure 3A). In 2 of the 5 Mtb/αCD4 NHP with reactivation, new granulomas that appeared during CD4 depletion had no viable Mtb growth (sterile) (Figure 3B). Thus, the similarity in necropsy score and bacterial burden observed between Mtb/αCD4 reactivators and non-reactivators is likely attributed to the fewer number of new granulomas and a low bacterial burden per granuloma during subclinical reactivation. In contrast, Mtb/SIV co-infected reactivators had substantial dissemination of Mtb resulting in new granuloma formation during reactivation that were more permissive to Mtb growth with higher bacterial burden.

**Figure 3.**
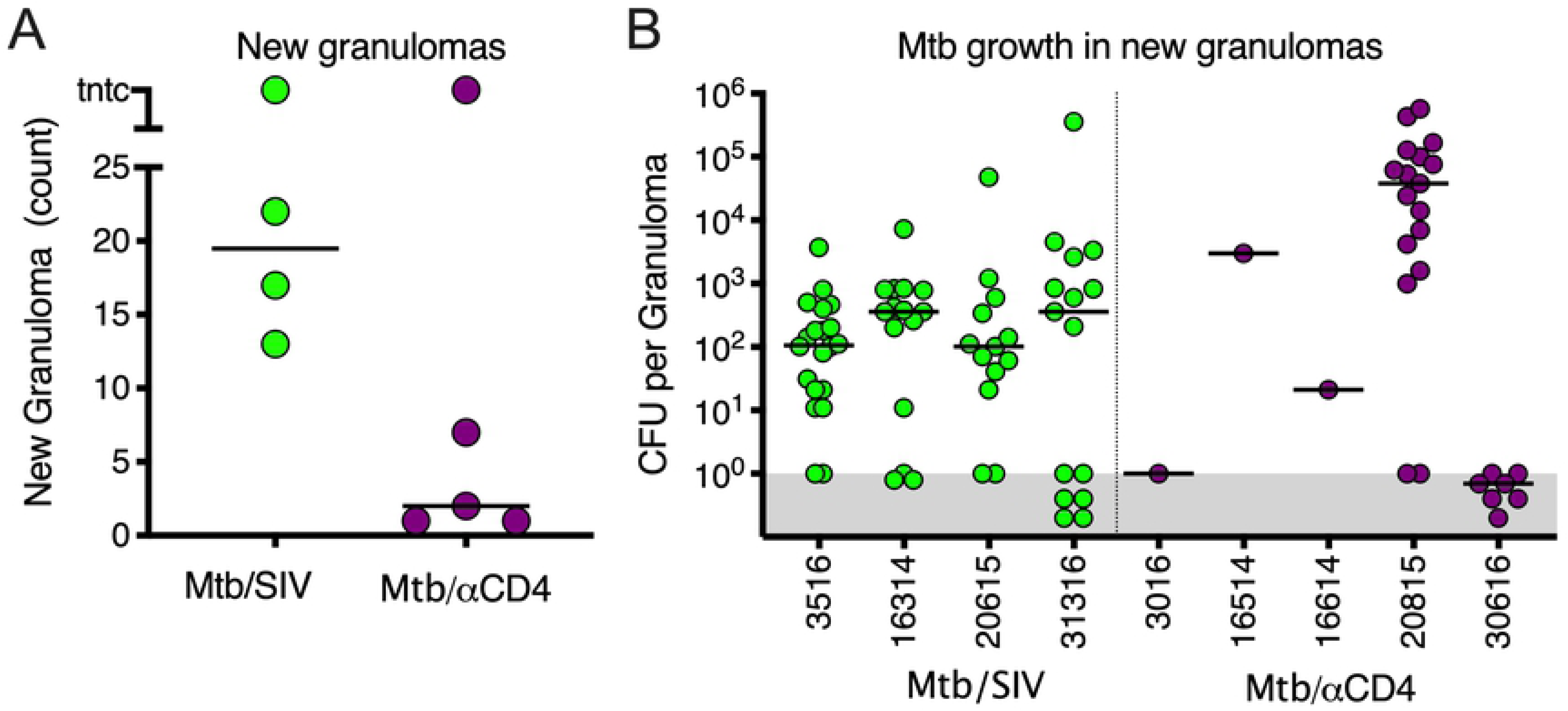
SIV-induced reactivation is characterized by more new granulomas that are permissive to Mtb growth compared to CD4 depletion. A) The number of newly formed granulomas identified by PET CT during subclinical reactivation, among Mtb/SIV (green, ranging from 13 to tntc) and Mtb/αCD4 (purple, ranging from 1 to tntc) NHP. TNTC = too numerous to count and was set at 100 (Mann-Whitney, P = 0.1270). B) Mtb growth from new granulomas of Mtb/SIV and Mtb/αCD4 NHP are shown. Points that fall within the grey bar were sterile. Numbers on x-axis represent individual monkey identification numbers. Lines represent medians. In A), each dot represents an individual animal; in B), each dot represents an individual granuloma.

Using barcoded Mtb strains matched with serial PET CT scans, we are able to track the dissemination of individual bacteria when there is Mtb growth [21]. We present one such case here. One of the Mtb/SIV co-infected animals had pre-existing granulomas in the right lower lung and developed new granulomas within the right lower and right middle lobe (Supplemental Figure 3) during subclinical reactivation. At least 3 of the 7 newly identified granulomas had barcodes that were also observed in the thoracic lymph nodes whereas the other new granulomas had similar barcodes detected in lung granulomas observed prior to SIV infection (Supplemental Figure 3). These data suggest that the Mtb dissemination during reactivation can occur from either the lung granulomas or thoracic lymph nodes. Interestingly, barcodes observed in extrapulmonary sites were similar to barcodes identified from lymph nodes in this animal, suggesting dissemination outside the lungs can occur from the lymph nodes.

### PET CT can predict subclinical reactivation from CD4 depletion but not Mtb/SIV co-infection

We assessed PET CT characteristics prior to immune suppression (SIV or αCD4) for the ability to distinguish reactivation risk, including total lung FDG activity, number of granulomas, greatest size or FDG avidity of any granuloma within an animal, and number of lobes involved (Supplemental Figure 4A-B). We did not observe any significant differences in PET CT characteristics prior to immune suppression that would distinguish reactivators from non-reactivators although the sample size was limited. We previously published that PET CT patterns during LTBI could discriminate macaques at high and low risk of TNF neutralization induced reactivation [18]. Specifically, total lung FDG activity (i.e., greater than 920 cumulative-SUV) and/or the presence of an extrapulmonary site of infection observed by PET CT prior to TNF neutralization predicted reactivation with 92% sensitivity and specificity [18]. By using these 2 metrics and the outcome of reactivation on animals in our current study, we could predict reactivation with 80% and 75% sensitivity and a specificity of 100% and 25% among Mtb/αCD4 and Mtb/SIV NHP, respectively (Supplemental Figure 4A). More specifically, the positive predictive value of high lung FDG activity and presence of extrapulmonary sites of infection was 100% among Mtb/αCD4 (i.e., high FDG activity and presence of extrapulmonary disease could predict reactivation 100% of the time) but only 50% of the Mtb/SIV animals. These data suggest that the spectrum of LTBI may predict reactivation risk that results from more specific immunologic impairments such as TNF neutralization or CD4 depletion. However, in the case of SIV infection where the immune suppression is broader, the threshold for reactivation to occur is less predictable.

### SIV and CD4 depletion modulate T cell composition within lung granulomas and thoracic lymph nodes

We quantified the T cells and assessed the quality of responses in thoracic lymph nodes and lung granulomas of both SIV and CD4 depleted animals (Figure 4, Supplemental Figure 5-9). Granulomas from Mtb/αCD4 NHP had significantly lower frequencies of CD4 T cells than those from either Mtb/SIV co-infected animals or latent Mtb-only controls (Figure 4A). CD8 T cell frequencies in granulomas from control Mtb-only animals were greater than both Mtb/SIV NHP and Mtb/αCD4 NHP (Figure 4A). We further examined differences in T cell frequencies between reactivators and non-reactivators in each experimental group (Figure 4B). Median frequencies of CD4 T cells were lower in granulomas from reactivated NHP in both the Mtb/SIV and Mtb/αCD4 groups compared to non-reactivators (Figure 4B). Interestingly, the absolute number (i.e., based on total number of cells estimated from the granuloma) of CD4 and CD8 T cells within granulomas from Mtb/SIV NHP was significantly greater than both latent control and Mtb/αCD4 animals (Figure 4C). Animals that developed reactivation in the Mtb/SIV group had greater absolute CD8 T cells though this apparently was not protective (Figure 4D). Absolute CD8 T cells were higher in Mtb/αCD4 NHP reactivators compared to non-reactivators (Figure 4D). The presence of greater absolute numbers of T cells in the granulomas during Mtb/SIV co-infection suggests that SIV infection alters the cellular composition and total quantity of T cells in the granulomas, although the increased T cells did not appear to improve disease outcome. This exemplifies a unique circumstance within the granuloma in which the frequency of T cells (i.e., the proportional contribution of T cells within an individual granuloma) may differ from the absolute number of T cells (i.e., the total contribution of T cells within the granuloma) between groups.

**Figure 4.**
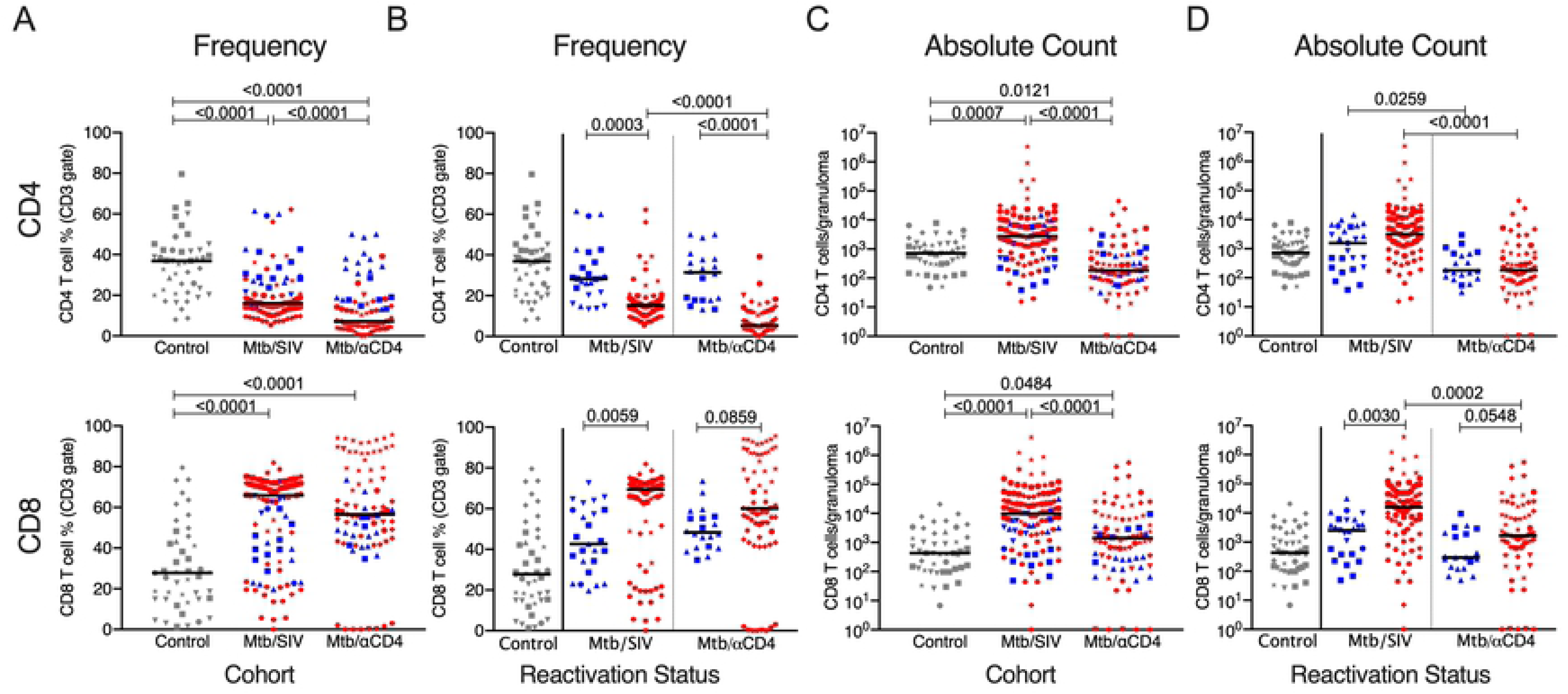
Frequencies of CD4 T cells are severely reduced by CD4 depletion but SIV markedly increases the total number of T cells within granulomas. Non-reactivators (blue) and reactivators (red) from Mtb only (control, grey), Mtb/SIV co-infected, and CD4 depletion (Mtb/αCD4) NHP are shown. A) and B) CD4 and CD8 T cell frequencies from lung granulomas (individual symbol) within individual monkeys (shapes) are shown. C) and D) Total number of CD4 and CD8 T cells within the granuloma are shown. Kruskal-Wallis test with Dunn’s multiple comparisons adjusted p-values are shown. P-values < 0.10 are shown. Lines represent medians. (6 Mtb control, n = 46 granulomas; 8 Mtb/SIV NHP, n = 110; and 7 Mtb/αCD4 NHP, n = 86; within Mtb/SIV NHP, non-reactivators = 25 granulomas, reactivators = 85; and within Mtb/αCD4 non- reactivators = 20, reactivators = 66).

Similar to lung granulomas, SIV infection and αCD4 antibody significantly reduced the frequency of CD4 T cells within thoracic lymph nodes compared to latent Mtb-only controls (Supplemental Figure 5A). Lower frequencies of CD4 T cells were observed in reactivators of Mtb/αCD4 NHP compared to non-reactivator (Supplemental Figure 5B), as previously published [5], although this pattern was not seen in the Mtb/SIV NHP. These data suggest that SIV infection influences thoracic lymph nodes in a different manner than αCD4 antibody and likely has important immunologic implications for reactivation.

### SIV and CD4 depletion change T cell cytokine production and granzyme B expression within lung granulomas

A homeostatic balance of pro- and anti-inflammatory responses that includes cytokine production and cytolytic function within granulomas is necessary for optimal control of Mtb [9]. To simplify the complexity of the functional immune markers data (cytolytic: granzyme B; T1/17 cytokines: TNF, IFN-γ, IL-2, IL-17; anti-viral: IFN-α; anti-inflammatory cytokines: IL-10 and IL-4) within granulomas, we used principal component analysis (PCA, Figure 5 & Supplemental Figure 6A-B). Principal component 1 (PC1) accounted for ∼ 60% of the variability for CD4 T cells; PC1 was characterized as T cell immune activation that includes IFN-α, IFN-γ, TNF, IL-2, IL-17, IL-10, IL-4 and granzyme B (Figure 5A and Supplemental Figure 6C). PC1 was similar for CD8 T cell responses in the granulomas, again accounting for over 60% of the variability of the data (Supplemental Figure 6D). The loading matrices for both cell types (Supplemental Figure 6A-B) show strong positive correlations of all of the cytokines with the component (ranging from 0.73 to 0.83 in CD4 cells and from 0.63 to in CD8 cells) suggesting that all functional markers (IFN-α, IFN-γ, granzyme B, TNF, IL-2, IL-4, IL-10, and IL-17) are driving the component uniformly. Among CD4 T cells, the median score of PC1 (i.e., a linear combination of functional immune markers) was greater among Mtb/SIV animals (regardless of reactivation status) compared to control and Mtb/αCD4 animals (Figure 5B). No difference was observed by CD4 T cells between Mtb/αCD4 and control animals (Figure 5B). Similarly, SIV granulomas had a greater CD8 median PC1 score compared to both Mtb/αCD4 and control groups (Figure 5B). Interestingly, the CD4 depleted groups had a higher CD8 T cell median PC1 score compared to controls. When PC1 scores were compared by reactivation status, greater scores were observed in granulomas from reactivated Mtb/SIV animals compared to those who did not reactivate. No difference between reactivation outcomes was observed in Mtb/αCD4 animals (Figure 5C). The immune parameters of PC1 in CD8 T cells also differentiated CD4 depletion induced reactivation from SIV infection. Of note, PC1 among CD4 and CD8 T cells was positively associated with Mtb burden (Figure 5D) and SIV replication (Figure 5E) within granulomas. Taken together, CD4 and CD8 T cells from Mtb/SIV granulomas have a more immune activated (more cytokines and granzyme B production) profile than Mtb/αCD4 and LTBI control groups though it is not protective. This pattern also correlates with reactivation status for Mtb/SIV NHP and increases with Mtb growth and SIV replication, suggesting that the pathogens and reactivation status correlate to homeostatic change in T cell activity within lung granulomas.

**Figure 5.**
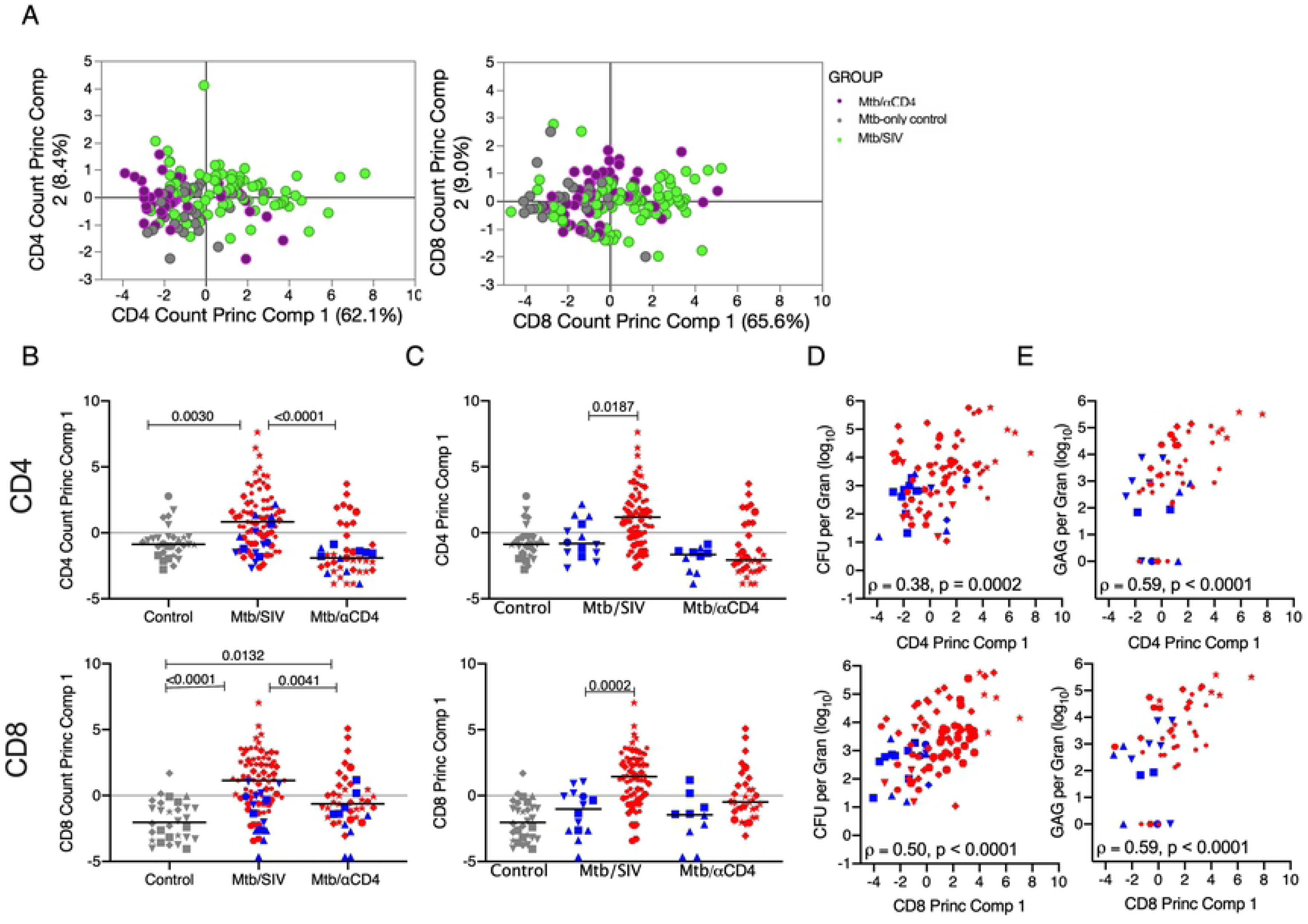
Principal component analysis demonstrates that SIV is associated with greater widespread T cells immune activation within the granuloma. A) Biplots are shown for the first two principal components. Each dot represents a granuloma. (Purple dots= granulomas from Mtb/αCD4 NHP, green dots = Mtb/SIV NHP, and grey dots = granulomas from Mtb-only control NHP. B) Median scores of of principal component 1 (includes total numbers of CD4 and CD8 T cells producing IFN-α, IFN-γ, TNF, IL-2, IL-17, IL-10, IL-4 and Granzyme B are compared across treatment groups. Non-reactivators (blue) and reactivators (red) from Mtb/SIV co-infected and Mtb/αCD4 NHP are shown. Individual monkeys are identified by different shapes. All treatment groups were compared using Kruskal-Wallis test with Dunn’s multiple comparison adjusted p-values reported. C) Median scores of principal component were compared between non-reactivators and reactivators within each treatment group. (Kruskal-Wallis Test with Dunn’s multiple comparison adjusted p-values reported.) D) The relationship between the CFU per granuloma (log_10_) and the first principal component was tested using Spearman’s ρ for all treatment groups. E) In Mtb/SIV NHP, the relationship between granuloma viral RNA quantification and the first principal component was tested using Spearman’s ρ. Each group contains the following number of granulomas: 30 Mtb-only Control, 43 Mtb/αCD4, 83 Mtb/SIV.

While PCA was used as a dimensional reduction method given the complex nature of the data sets, we also performed more traditional analytic methods comparing groups by single immune functional parameters with results that were generally consistent with the PCA results (Supplemental Figure 7). Lung granulomas from Mtb/SIV NHP also contained more (p = 0.0528) CD38+ T cells compared to Mtb-only granulomas, suggesting that the Mtb/SIV co-infection is associated with increased T cell activation (Supplemental Figure 8). SIV co-infection and CD4 depletion also changed the overall composition of T cells within lung granulomas that produce these cytokines and granzyme B (Supplemental Figure 9). Surprisingly, non-traditional CD3 T cells (i.e., CD3+CD4-CD8- T cells, and CD3+CD4+CD8+ T cells) actively contributed to the overall immune function of these granulomas. CD3+CD4+CD8+ T cells have distinct cytokine and cytolytic responses within PBMC, BAL, and lung granulomas compared to traditional CD4 and CD8 T cells within *M. tuberculosis* infected cynomolgus macaques [27]. The responses from these non-traditional T cells suggest that SIV and CD4 depletion disrupt both conventional and non-conventional T cells types within the granuloma. Overall, these data suggest that SIV co-infection causes a dysfunctional T cell homeostatic function and pro/anti-inflammatory balance that differs from CD4 depletion.

### Mtb increases SIV replication and SIV replication reduces Mtb killing within the granuloma

The ability of Mtb to increase HIV replication has been demonstrated *in vitro* under specific conditions [30, 31]. Plasma viremia during the course of acute infection was compared between Mtb/SIV animals and SIV only control animals. Viremia was significantly higher at 1-week post-SIV infection in the Mtb/SIV animals, although viremia reached a similar level in both groups by 2 weeks (Figure 6A). Overall, similar PBMC CD4 and CD8 T cell frequencies and numbers were observed in Mtb/SIV and SIV-only NHP (Supplemental Figure 10).

**Figure 6.**
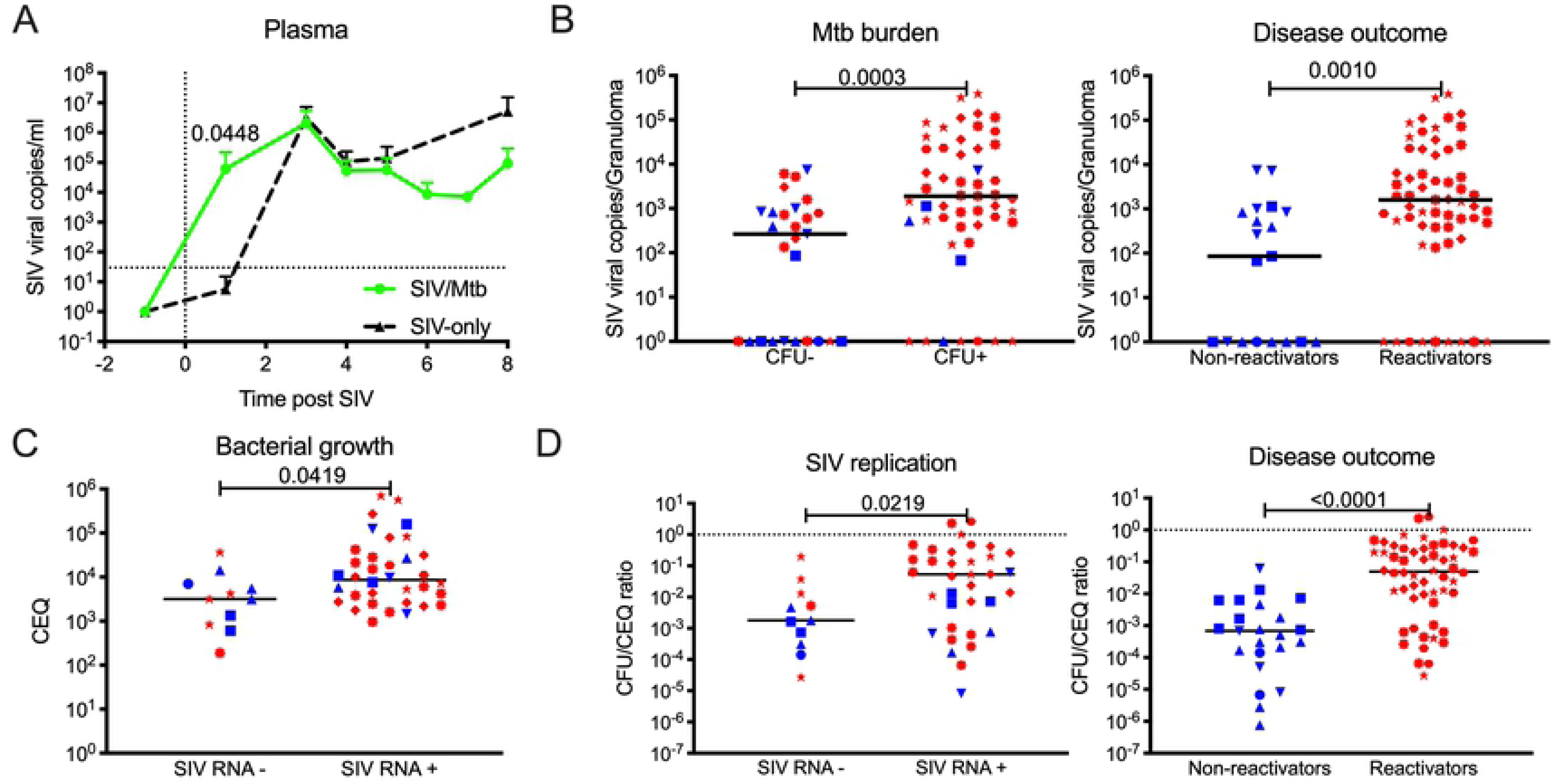
SIV replication within the granuloma is associated with reactivation status, greater bacterial burden and growth with less bacterial killing. A) Comparison between plasma SIV RNA copies/ml from SIV-only (black, n = 4) and Mtb/SIV co-infected (green, n = 8) NHP. Symbols represent means and error bars represent standard deviations. B) Differences in SIV RNA replication within lung granulomas from Mtb/SIV co- infected NHP by Mtb burden (Mtb growth [CFU+, n = 51] and sterile [CFU-, n = 27]) and outcome (reactivators [red, n = 59] and non-reactivators [blue, n = 19]). Each symbol is a granuloma. C) Bacterial growth (presented as chromosomal equivalents, CEQ) is greater within among Mtb/SIV granulomas with detectable SIV RNA (SIV-, n = 11; SIV+, n = 35). D) Within Mtb/SIV NHP granulomas, less bacterial killing (represented as CFU/CEQ ratio) is observed when SIV RNA is present (SIV-, n = 11; SIV+, n = 35) and during reactivation (reactivators [red, n = 56] and non-reactivators [blue, n =23]). CFU was transformed by adding 1 to reflect sterile lung granulomas with CEQ. Dotted line at Y = 1 defines no killing. Two points above Y = 1 represent the higher CEQ threshold (1000) compared to CFU’s lower threshold (10). Each shape represents an individual NHP. Individual t tests were utilized to determine significant differences (P < 0.05) between SIV-only and Mtb/SIV NHP. The Mann-Whitney test was used to determine significance between groups in granulomas. Lines represent medians in B-D.

HIV infection has been detected in TB diseased lungs [32], lymph nodes [33], pleural fluid [34], and cerebral spinal fluid [35]. While SIV has been identified within lung and lymph nodes of Mtb/SIV co-infected NHP [11, 15], limited data exist regarding SIV or HIV infection in individual Mtb granulomas [10]. To address this, we examined the level of cell-associated SIV RNA within individually harvested granulomas from Mtb/SIV co-infected animals and found that CFU+ granulomas (measured as Mtb colony forming units, CFU) were associated with higher SIV RNA copy numbers (Figure 6B). When sorted by outcome, granulomas from animals that reactivated compared to those that did not reactivate had higher SIV RNA copies (Figure 6B), consistent with the positive correlation between SIV replication and Mtb burden in lung granulomas (Supplemental Figure 11A). To ensure that the increased SIV RNA copies per granulomas were not simply due to increased numbers of CD4 T cells in Mtb/SIV animals, we compared SIV RNA:CD4 RNA ratios (Supplemental Figure 11B). Higher RNA SIV:CD4 RNA ratios were associated with granulomas from reactivated animals compared to non-reactivators and in granulomas with Mtb burden (Supplemental Figure 11B).

To determine whether SIV alters the ability of a host to kill Mtb *in vivo*, we compared ratios of live Mtb (CFU) to total (dead and live) bacteria (measured as chromosomal equivalents; CEQ) as an estimate of bacterial killing [19, 36]. Lung granulomas with SIV RNA contain higher CEQ than lung granulomas without SIV RNA (Figure 6C), indicating increased bacterial growth. Here, lower CFU/CEQ ratios indicate more Mtb killing whereas higher ratios indicate poor killing. Granulomas from Mtb/SIV NHP reactivators had reduced Mtb killing (higher CFU/CEQ) compared to those from non-reactivators (Figure 6D). Similarly, reduced Mtb killing was observed in granulomas with detectable SIV RNA. Taken together, these data support that there is synergy between local Mtb and SIV replication dynamics *in vivo* that coincides with increased Mtb growth, reduced Mtb killing and increased virus replication within the granuloma itself.

## Discussion

Here, we are able to identify the early dynamics of reactivation from LTBI by focusing on the events that occur during subclinical reactivation. In this highly controlled experimental setting, both CD4 depletion and SIV infection after established LTBI induced subclinical reactivation but with different immunologic mechanisms and patterns of reactivation. It should be noted that SIV_mac251_ preferentially infects and depletes CCR5+ CD4 T cells (especially CD45Ra-memory cells) in the periphery [37], while CD4 depletion antibody causes a broader depletion of CD4 T cells. In this study, SIV infection induced only a transient decrease (3 weeks post-SIV infection) in the total peripheral CD4 T cell frequency but otherwise was similar to control groups and no differences were noted in the absolute number of circulating CD4 T cells in blood allowing us to compare the CD4 dependent and independent mechanisms of reactivation. Notably, quantification of CD4 T cells in blood did not consistently reflect CD4 T cells in the granulomas as was the case for Mtb/SIV co-infection groups. The use of PET CT allows us to capture the earliest events of reactivation during immune suppression before overwhelming bacterial burden, disease pathologies and overt clinical signs develop. Both SIV infection and CD4 depletion during LTBI induced similar rates of subclinical reactivation, based on the strict definition of detection of a new granuloma by PET CT (and confirmed at necropsy), but the bacterial burden and severity of dissemination was worse in the Mtb/SIV group. This was attributed to the greater number of new granulomas that developed with viable Mtb growth among Mtb/SIV NHP despite having significantly more CD4 T cells than Mtb/αCD4 group. Importantly, CD4 depleted animals had fewer new granulomas during early reactivation and many were sterile suggesting that CD4 independent mechanisms for Mtb killing exist within the granuloma during LTBI as others have suggested [15]. Within the granuloma, immune activation profiles were more perturbed by SIV than by CD4 depletion and the presence of SIV RNA copies was associated with greater bacterial growth and reduced bacterial killing.

Understanding the mechanisms of HIV-induced reactivation of LTBI is a critical question that has been the focus of NHP studies. Recently Bucsan et al. demonstrated that CD4 depletion resulted in reactivation in only 1 out of 8 latently infected rhesus macaques based on overt clinical signs in contrast to their historical data in which 9 of 17 LTBI animals developed reactivation after SIV_mac239_ [38]. These differences in outcome may be attributed to inherent differences in the macaque models and Mtb strain used for infection. In their rhesus macaque model of Mtb infection, LTBI is established 9 weeks after inoculation with low virulence Mtb CDC1551 strain [39, 40] and CD4 depleting antibody was given for up to 9 weeks. In our previous reactivation studies in our cynomolgus macaque model of LTBI using a low dose virulent strain of Mtb, 50% reactivation was observed in animals that underwent CD4 depletion [5] while 100% of animals infected with SIV_mac251_ reactivated, although only half occurred by 8 weeks after SIV infection [11]. Our rates of subclinical reactivation defined by PET CT in our current study are consistent with our previously published data. Importantly, we were able to use subclinical reactivation (appearance of a new granuloma) as our endpoint rather than overt clinical reactivation, taking advantage of the fact that PET CT facilitates a more in-depth understanding in the pathogenesis of the earliest phases of reactivation. We were able to identify by PET CT the new granulomas that emerged during CD4 depletion and harvest them at necropsy; surprisingly, no viable Mtb could be recovered from a subset of these. This reinforces the notion that granulomas can sterilize despite having few to no CD4 T cells and highlights the complex, heterogeneous nature of granuloma function in which CD4 T cells may not be required by all granulomas to contain Mtb growth. This is consistent with our prior reactivation studies in which the a subset of newly developed granulomas had no culturable Mtb during reactivation after TNF neutralization [18]. These data further characterize the early events of reactivation during LTBI.

The immunologic mechanisms of Mtb susceptibility among HIV-infected individuals remain unclear given the unknown timing and order of HIV or Mtb infection, highly variable nature of clinical studies, and limited access to tissue samples in humans [10]. We did not find overt PBMC immune responses that correlated with reactivation in either the Mtb/αCD4 or Mtb/SIV groups. HIV/Mtb co-infection clinical studies have demonstrated that HIV preferentially induces an overall reduction in Mtb-specific IL-2 producing peripheral CD4 T cells without changes in cytomegalovirus-specific CD4 T cell responses [41], Mtb-specific Th1 producing peripheral T cells [42, 43], broad spectrum of Th transcription factors within Mtb-specific CD4 T cells [44], and PPD (purified protein derivative from Mtb)-stimulated BAL CD4 T cells producing IFN-γ and IL-2 [43]. Reductions in the frequency of CD4 and CD8 T cell production of Type 1 and IL-17 cytokines have also been observed in the blood of co-infected patients [45, 46]. In one study, 5 patients with newly acquired HIV infection and LTBI were found to have reduced frequencies of Mtb-specific CD4 T cells though none developed reactivation in the first year of seroconversion [41]. Thus, HIV directly dampens Mtb-specific T cell functions that are essential for control of Mtb infection within blood and BAL, which highlight the need for studies that examine immunologic changes within lung granulomas. These data are consistent with the increased rates of TB reported during the first year of HIV seroconversion despite normal CD4 T cell counts [47]. HIV or SIV may cause Mtb-specific CD4 T cells in blood to migrate to the lungs in response to Mtb infection, but without tetramers to identify Mtb-specific T cells we cannot be certain, which is a limitation of this study. As HIV infection progresses, severe depletion of peripheral CD4 T cells correlates with increased Mtb presence within granulomas of HIV/Mtb co-infected individuals [10] and loss of interstitial CD4 T cells with an increase in Mtb growth in co-infected humanized mice [14]. It is important to understand how HIV directly affects granulomas as responses in PBMC, BAL, and granulomas are not well correlated, as observed in this study and others [11, 12, 14, 15, 43].

Granulomas have been hypothesized to be ideal sites for HIV replication [reviewed in [10, 48–50]]. We quantified individually harvested granulomas from SIV-Mtb co-infected animals and observed more SIV RNA among reactivators, suggesting that SIV replication within granulomas is directly associated with disease status. These data are consistent with findings from others in which more SIV-infected cells within lung tissue were observed in Mtb/SIV reactivators compared to non-reactivators by immunohistochemical analysis [15], although that study did not examine individual granulomas. Similarly, more SIV DNA was obtained from lung tissue of active Mtb/SIV co-infected rhesus compared to latently infected rhesus macaques [16], indicative of more infected cells in co-infected lungs. Importantly, we show that SIV transcription was significantly higher in granulomas with live Mtb, which was associated with reduced Mtb killing. This is consistent with *in vitro* studies showing that HIV infection reduces TNF-mediated macrophage apoptosis [51, 52] and that the presence of culture filtrate protein from Mtb is associated with increased cellular HIV replication [53]. To the best of our knowledge, our data represent the largest collection of granuloma-specific quantification of Mtb and SIV RNA in the literature to date and underscore the important synergy between these two pathogens directly within the granuloma.

The factors that influence this synergy between Mtb and SIV have been hypothesized for years, and assessed peripherally, in airways and *in vitro*, but not in a realistic *in vivo* tissue-based setting such as lung granulomas. Although the precise mechanisms for increased SIV RNA and reduced Mtb killing in individual granulomas were not identified in this study, we hypothesize that HIV induces a combination of mechanistic immunological changes within lung granulomas that result in Mtb growth and disease progression (Figure 7). Elucidating how HIV or SIV manipulates the micro-environment of Mtb granulomas is a critical factor in understanding the mechanisms of HIV-Mtb co-infection and subsequent treatment and prevention. Lung granulomas are heterogeneous with variable T cell composition, functions and Mtb growth or killing (and SIV during co-infection) within a single host [9, 12]. Within the granuloma, SIV infection significantly altered the composition of T cells and their function by increasing the overall production of both pro- and anti-inflammatory cytokines and granzyme B by CD4 and CD8 T cells. This increase in activation was associated with reactivators and correlated with Mtb growth and SIV replication. We also observed a greater frequency of activated (CD38+) T cells within granulomas from SIV-Mtb infected animals compared to LTBI control [54]. Despite the increase in T cell activation and effector production in granulomas, Mtb/SIV NHP still experienced reactivation of LTBI, increased Mtb growth, and reduced ability to kill Mtb within granulomas. This suggests that the increase in activation and function of T cells may be in response to SIV-induced earlier loss of control of Mtb infection, however this response is too late to restrain Mtb. Further studies at earlier time points post-SIV infection in this model are necessary to determine the early effects of SIV on the granuloma environment.

**Figure 7.**
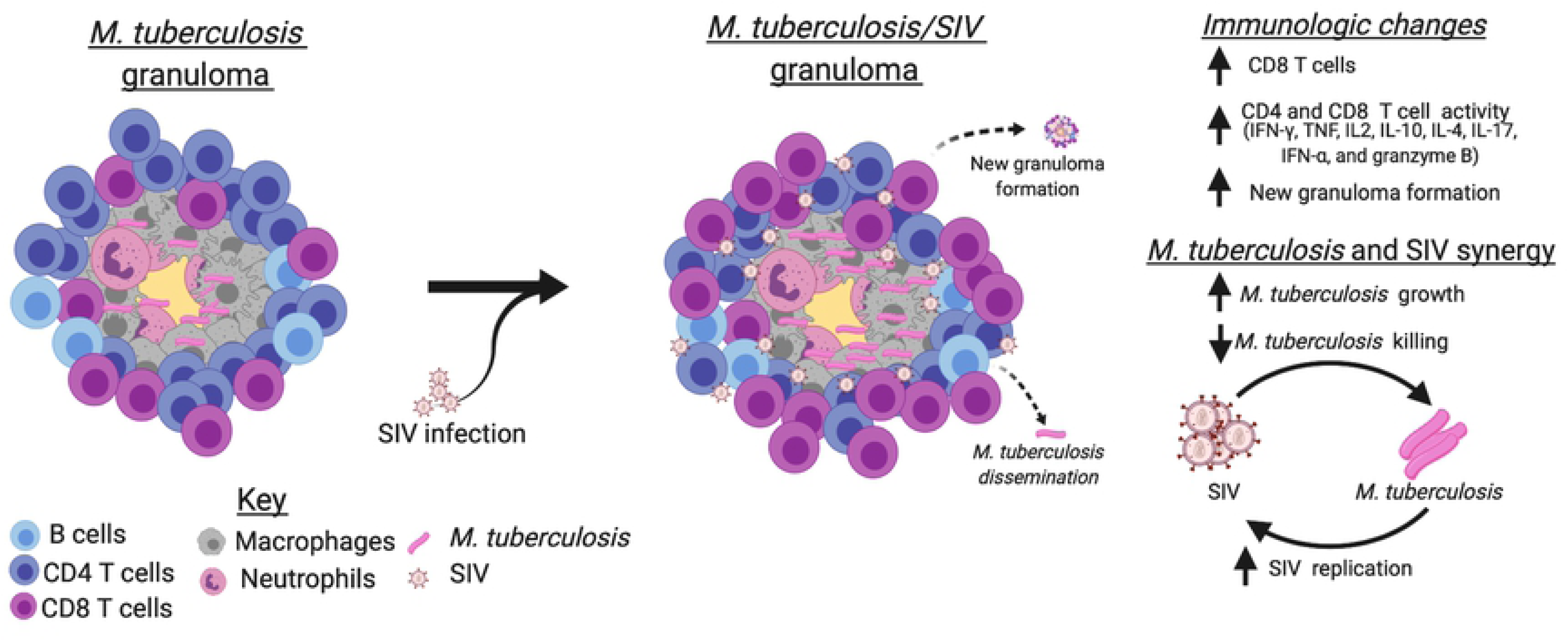
SIV changes immunological functions within Mtb lung granulomas, increases Mtb growth, and reduces Mtb killing. A) An example Mtb caseous granuloma contains T cells, macrophages, neutrophils, and Mtb. B) Mtb/SIV co-infected granulomas contain more CD8 T cells and an increase in overall production of Th1 cytokines, granzyme B, IL-17, IL-10, IL-4, and IFN-α by CD4 and CD8 T cells. SIV also increases the probability of causing new granulomas to form, Mtb growth and dissemination. SIV has been linked to increases in Mtb growth and a reduction in Mtb killing during reactivated disease, while Mtb growth correlates to increases in SIV replication. This suggests that lung granulomas are sites that support a synergistic relationship between SIV replication and Mtb growth. Image created by BioRender.com.

Interestingly, we observed increased Type1, Type17 and IL-10 cytokine producing CD8 T cells within Mtb/SIV reactivators, which is important because IFN-γ, TNF and IL-17 are necessary for activating macrophages to kill Mtb [55, 56]. Greater production of granzyme B (predominantly from CD8 T cells) was observed in the Mtb/SIV NHP that reactivated compared to those that did not. This is in contrast to other reports in which CD8 production of granzyme B was associated with protection from reactivation [15]. Although granzyme B is a cytolytic molecule that can induce apoptosis in infected macrophages and kill Mtb directly [57], HIV has been reported to impair degranulation of cytoplasmic granules, such as granzyme B [58]. In this study, more CD4 and CD8 T cells expressing IFN-α and IL-10 resided in lung granulomas from Mtb/SIV NHP that reactivated in this study. Highlighting the complexity of cytokine production within granulomas, in another study Mtb growth was correlated to IFN-α and IL-10 production within cervical lymph nodes with Mtb granulomas from anti-viral treatment naïve HIV/Mtb co-infected patients [59], which suggest that these cytokines might play a role in regulating Mtb growth or reflect generalized immune activation induced by SIV. A limitation of our study is that we did not examine macrophage functions within lung granulomas. We hypothesize that HIV is disrupting the interaction between macrophages and T cells within granulomas, so future studies will examine the complex macrophage environment of the granuloma in more detail.

Given the complex nature of the immune cells and the heterogeneous function of granulomas during Mtb infection, the optimal protective function of the granuloma is likely not dependent on a single immune mechanism or cell type but rather on a combination of possible immune interactions within the granuloma (reviewed in [60]), particularly as the increase in total T cells and increased activation of T cells within granulomas was not protective. While we focused on the role of T cells in this study, macrophages certainly play a key role in Mtb killing and in SIV infection in addition to other immune cells within the granuloma that we did not directly assess, which is an important limitation in this study. While no single cell type or immune mechanism was associated with reactivation in our studies, we hypothesize that the overall balance of both pro- and anti-inflammatory properties necessary for optimal granuloma function [9, 60] is disrupted by the complex nature of SIV infection compared to CD4 depletion alone, accounting for the more dramatic reactivation pattern in Mtb/SIV macaques. Given the heterogeneous nature of individual granulomas, any perturbation that leads to a more pro-inflammatory or more anti-inflammatory state could be permissive for both Mtb (reviewed in [61]) and SIV replication [62]. However, our data suggest that many granulomas are able to contain Mtb independent of CD4 T cells, which was also true in TNF-neutralized macaques [18]. Clearly the pleiotropic immune perturbations from SIV or HIV infection within the granuloma lower the threshold for reactivation in a multifactorial and dramatic fashion. Thus, strategies in vaccination or host-directed therapy for HIV-Mtb co-infected individuals are likely to require a multifactorial approach given the complex nature of granulomas.

## Author contributions

CRD and PLL wrote manuscript. CRD, ZA, SF, JLF, PLL developed experiments. CRD, TR, TMB, PM, PK, AGW, HJB, FH, JB performed experiments. Statistical analysis performed by PM, CRD, PLL. All authors edited and commented on manuscript.

## Acknowledgements

We thank the tireless efforts of our veterinary technicians/imaging staff (Melanie O’Malley, Jaime Tomko, Daniel Fillmore, Chelsea Causgrove, Brianne Stein, L. James Frye) and research technicians (Cassaundra Ameel, Nicholas Schindler, Carolyn Bigbee, Amy Myers, Mark Rodgers, Catherine Cochran, Chris Kline). Special thanks to Charles Scanga for study coordination and members of the Flynn, Mattila, Gideon, and Scanga labs for their helpful discussion. These studies were funded by the National Institutes of Health, National Institutes of Allergy and Infectious Diseases R01 AI11871 (PLL), AI134195 (PLL), and Otis Childs Trust of the Children’s Hospital of Pittsburgh Foundation (PLL). CD4 depleting antibody was produced by the NIH Non-Human Primate Reagent Resource (R24 OD010976, U24 AI126683). SIV gag/pol peptides were obtained from the NIH AIDS Reagent Program, Division AIDS, NIAID. The authors have declared that no conflict of interest exists.

## Supplemental Data Captions

**Supplemental Table 1.** Antibodies used for intracellular cytokine staining

**Supplemental Table 2.** Clinical signs and disease outcome for each nonhuman primate in study

**Supplemental Figure 1. Example gating strategy for flow cytometry.** A) Singlet events positively selected. B) Live cells negatively selected. C) Lymphocytes selected. D) CD3 T cells positively selected and CD4 and CD8 T cells selected from CD3 T cell gate. E) Example cytokine and granzyme B expressing T cells. TNF, IFN-γ, and granzyme B are displayed. Gating example from a lung granuloma.

**Supplemental figure 2. Extrapulmonary disease (EP) and Mtb growth within thoracic lymph nodes were positively correlated within Mtb/SIV NHP.** Correlation between EP score and Lymph node Mtb growth within control, Mtb/αCD4 and Mtb/SIV was compared. Spearman’s test was performed and correlation coefficient (rho) and p values < 0.05 are presented. Each dot represents an animal. NHPs that reactivated are represented in red and non-reactivators are represented in blue.

**Supplemental Figure 3. Tracking of Mtb lesions with barcodes over time.** The panel on the left shows lung granulomas (small dots) and lymph nodes (large pie charts) that were seen prior to SIV infection on PET-CT scans. The right panel shows the barcodes from granulomas and lymph nodes seen only post-SIV infection (in color) with the barcodes identified within granulomas prior to SIV infection shown in black. Extrapulmonary barcodes are shown below the lung renderings. All extrapulmonary tissues represented here were identified only after SIV-infection at necropsy. Colors denote barcode content. Solid colors indicate a sample which contained only one barcode, while pie chart markers reflect the relative barcode content of samples which contained two or more barcodes.

**Supplemental Figure 4. PET CT characteristics prior to immune suppression do not predict reactivation in either SIV or αCD4 antibody treated animals.** Each dot represents an individual animal. A) Total lung FDG activity prior to SIV infection or αCD4 depletion (dotted line set at the TNF-induced predictive reactivation threshold value) is shown among reactivators (red) and non-reactivators (blue). Open circles represent animals with extrapulmonary disease evident on scan before immune suppressant. B) FDG uptake per granuloma, number of lung lobes containing granulomas, total granuloma counts, and size (in mm) of largest granuloma are compared between reactivators and non-reactivators. Kruskal-Wallis performed, all p-values > 0.10; therefore none are reported. TNTC = too numerous to count.

**Supplemental Figure 5.** C**D**4 **T cell frequencies are reduced within thoracic lymph nodes of Mtb/SIV and Mtb/αCD4 NHP**. T cell frequencies and total counts from thoracic lymph nodes (individual symbols) within individual monkeys (shapes) from non-reactivators (blue) and reactivators (red) and controls (grey). A) Differences in CD4 and CD8 T cell presence within infection cohort (Mtb only, control, n = 27; Mtb/SIV, n = 40; and Mtb/αCD4, n = 27) are presented. B) Differences in CD4 and CD8 T cell presence based on disease outcome (reactivator; non-reactivator) are presented. Within Mtb/SIV NHP, non-reactivators = 21 thoracic lymph nodes, reactivators = 19; and within Mtb/αCD4 non-reactivators = 8, reactivators = 17. Lymph nodes with granulomas are represented by large symbols and the small symbols identify lymph nodes without granulomas. P values reported represent Kruskal-Wallis test with Dunn’s adjusted p-values are show P-values < 0.10 are shown. Lines represent medians.

**Supplemental Figure 6. Results of Principal Component Analysis on CD4 and CD8 cytokine counts.** Biplots of the first two principal components on CD4 (A) and CD8 (B) counts. For both CD4 and CD8 counts, the first principal component represents over 60% of total variability of the entire sample of granulomas. The loading matrix displays the correlation of each individual cytokine with the principal component for CD4 T cells (C) and CD8 T cells (D). In CD4 counts, IFN-α has the strongest correlation with the component (0.83264); in CD8 counts, IFN-γ has the strongest correlation (0.87519). Each group contain the following number of granulomas: 30 Control, 43 aCD4/Mtb, 83 SIV/Mtb.

**Supplemental Figure 7. SIV changes CD4 and CD8 T cell cytokine and granzyme B expression within lung granulomas compared to Mtb-only NHP** Absolute counts of cytokine production and granzyme B presence within CD4 and CD8 T cells of lung granulomas from Mtb-only (grey symbols), Mtb/SIV, and Mtb/αCD4 from NHP and from non-reactivated (blue) and reactivated (red) NHP. Each symbol is a lung granuloma and individual NHP are represented as different shapes. Kruskal-Wallis with Dunn’s adjusted p-values are reported, accounting for the following (4) comparisons: reactivator vs non-reactivator within each group and reactivators and non-reactivators across groups (Reactivators: Mtb/SIV vs Mtb/αCD4, non-reactivators: Mtb/SIV vs Mtb/αCD4). P-values < 0.10 are shown. Lines represent medians. The number of granulomas within each group are as follows-Cytokine and Th1 cells (100 CD3 T cell threshold): 6 Mtb only, n = 30; 8 Mtb/SIV, n = 83; and 7 Mtb/αCD4 NHP, n = 43; Mtb/SIV 4 reactivators, n = 69, 4 non-reactivators, n = 14; Mtb/αCD4 NHP, 5 reactivators, n = 33, 2 non-reactivators, n = 10).

**Supplemental Figure 8. More activated T cells are within lung granulomas of Mtb/SIV compared to Mtb-only NHP.** A) Immunohistochemistry images of nuclei (blue), CD38 (green), CD3 (red) images from Mtb-only and Mtb/SIV NHP lung granulomas. Arrows identify CD3+CD38+ T cells. B) CD38+CD3+ T cells were quantified from 6 Mtb/SIV (n = 13) and 6 Mtb-only (n = 11) NHPs. Reactivators are identified in red and non-reactivators in blue. Each symbol represents a granuloma and each shape represents a different NHP. Quantification was performed on regions of interest (ROI, 20x image) of lung granulomas. Mann-Whitney test was used to determine significance between groups in granulomas (p value displayed). Lines represent median.

**Supplemental Figure 9. Changes in T cell composition of Mtb-specific cytokines and cytolytic markers during SIV infection and CD4 depletion within the granuloma.** The distribution of T cells within the granuloma are represented on the left panels represented as CD4 T cells (CD4+CD8-, green), CD8 T cells (CD8+CD4-, orange), other T cells (CD4+CD8+ T cells, grey and CD4-CD8-T cells, black). The distribution of T cells making any Mtb-specific cytokines or cytolytic markers is shown to the right. Permutation tests were used to compare groups. The number of granulomas within each group are as follows-CD3 T cells: Mtb only, n = 47; Mtb/SIV, n = 110; and Mtb/αCD4 NHP, n = 86. Cytokine and Th1 cells (100 CD3 T cell threshold): 6 Mtb only, n = 30; 8 Mtb/SIV, n = 83; and 7 Mtb/αCD4 NHP, n = 43; Mtb/SIV 4 reactivators, n = 69, 4 non-reactivators, n = 14; Mtb/αCD4 NHP, 5 reactivators, n = 33, 2 non-reactivators, n = 10).

**Supplemental Figure 10. Changes in peripheral blood mononuclear cells (PBMC), bronchoalveolar lavage (BAL), and peripheral lymph node (pLN) T cells within Mtb/SIV and SIV-only NHP.** Changes in CD4 and CD8 T cell counts (Abs Counts) and frequencies after SIV_mac251_ infection (green line, Mtb/SIV, n = 8) or SIV-only (black line, n = 4). Statistics reported are Wilcoxon-exact tests comparing Mtb/SIV and SIV-only at each time point (not adjusted for multiple tests). Lines are median and error bars represent interquartile range. **** p < 0.0001, *** p < 0.001, ** p < 0.01, * p < 0.05, # p < 0.10.

**Supplemental Figure 11. SIV replication correlates to Mtb growth and is not attributed higher levels of CD4 T cells alone in the granuloma.** A) A positive correlation between lung granulomas that contain both SIV replication and Mtb growth (n = 42) is observed. B) Granuloma specific ratios of SIV viral RNA (vRNA): CD4 RNA is shown among SIV/Mtb animals who were non-reactivators (n = 22) and reactivators (n = 58) as well as from granulomas with Mtb growth (CFU+, n = 52) and without viable Mtb growth (CFU-, n = 28). Each symbol is a granuloma and individual NHP are represented as different shapes. Red symbols identify reactivators and blue symbols identify non-reactivators. Samples without SIV replication or Mtb growth are presented as a reference. All data was log10 transformed. Pearson correlation coefficients are reported with corresponding p-values in A. The Mann-Whitney test was used to determine significance between groups in granulomas. Lines represent medians in B.

